# Local niche-derived immunosuppressive CXCR2^+^ cells impair antiviral immunity

**DOI:** 10.1101/2025.08.24.671975

**Authors:** Akisawa Satomi, Riho Saito, Tadahaya Mizuno, Hiroki Sugishita, Hideki Ukai, Shigeyuki Shichino, Masashi Yanagisawa, Kouji Matsushima, Yukiko Gotoh, Tomohiko Okazaki

## Abstract

Symptom severity after viral infection varies among individuals, yet its mechanism remains poorly understood. We categorized mice into recovering and non-recovering groups based on body weight after infection of the same titer of vesicular stomatitis virus (VSV). We revealed that in the olfactory bulb (OB) where VSV initially expands, non-recovering mice exhibited an anti-inflammatory environment, suggesting its detrimental effect. Importantly, CXCR2^+^ cells resembling immunosuppressive myeloid-derived suppressor cells (MDSCs) were more abundant in the OB of non-recovering mice than in that of recovering mice after VSV infection. Depleting CXCR2^+^ MDSC-like cells from the brain increased inflammatory responses and the animal’s survival after infection. Furthermore, site-specific labeling indicated that a significant fraction of these cells in the OB originate from the skull-bone marrow (skull-BM) as well as circulation. This study reveals the lethal effects of CXCR2^+^ MDSC-like cells on local immune responses to viral infection, highlighting their therapeutic potential for antiviral defense.

## Introduction

Viral infections lead to a wide range of pathological outcomes in infected individuals, from mild to severe^1–3^. Understanding the mechanisms underlying these variations is challenging because the manifestation of pathological outcomes is heavily influenced by the genetic background of the infected host, as well as environmental factors (e.g., infection history)^3–5^. Intriguingly, the pathological severity following viral infection can differ between individuals that share similar backgrounds in animal models^6–8^. For example, intranasal infection of mice with the same titer of vesicular stomatitis virus (VSV) results in death of some animals but not others, even under apparently identical genetic and environmental conditions^6,7^. These findings suggest the existence of an as-yet-unidentified mechanism that governs life-or-death outcomes.

What could be the causes leading to the severe outcome of an infected host? One is functional impairment of infected tissues and organs triggered by viral replication. During this process, viruses induce direct cytopathic effects or produce viral virulence factors that damage host tissues^9^. Host immune responses mitigate such virus-induced pathology by restricting viral replication through the upregulation of antiviral and inflammatory genes^10^. On the other hand, excessive activation of host immune responses can also disrupt tissue homeostasis and lead to more severe disease outcomes^11,12^. For instance, severe influenza virus or SARS-CoV-2 infections are often characterized by cytokine storms, which are associated with high mortality^13–15^. Thus, antiviral immune responses represent a double-edged sword: when induced with appropriate timing and magnitude, they enable both pathogen clearance and the maintenance of tissue homeostasis, thereby contributing to host survival.

These magnitude and timing of immune responses are critically shaped by the interplay between pro-inflammatory effector cells and immunoregulatory populations. Innate immune cells such as natural killer (NK) cells, neutrophils, monocytes, macrophages, and dendritic cells constitute the first line of defense by rapidly responding to infection through pathogen sensing, cytokine production, and direct effector functions such as cytotoxicity or phagocytosis^16^. These responses are complemented by adaptive immune cells, including cytotoxic CD8^+^ T cells and CD4^+^ helper T cells, which provide antigen-specific immunity through targeted killing of infected cells and by supporting antibody production and enhancing cellular immune responses^16^. In parallel, immunosuppressive populations act to prevent immune-mediated tissue damage, with cells such as regulatory T cells (Tregs) and myeloid-derived suppressor cells (MDSCs) contributing to immune suppression through multiple mechanisms, including the production of anti-inflammatory cytokines such as IL-10 and TGF-β, and the expression of immune checkpoint molecules ^17,18^. Understanding balancing mechanism for these opposing forces is essential for clarifying how the immune system navigates the trade-off between effective viral control and the preservation of tissue integrity.

The functional interplay among various immune cells within infected tissues raises another question: where do these immune cells originate, and how are they supplied to sites of infection? In peripheral tissues, resident immune cells contribute to homeostatic regulation and baseline immune surveillance. Upon infection, however, additional immune cells are rapidly recruited from the circulation to initiate robust antiviral responses at the site of infection. Beyond these two well-characterized sources— resident cells and blood-derived populations—emerging evidence highlights a third, distinct source of immune cells: those derived from microenvironments that are directly connected or physically adjacent to the target tissue^19^. These local immune niches can supply immune cells independently of vascular trafficking and contribute to shaping timely and spatially confined responses^20,21^. Together, these immune cell populations derived from diverse sources act in concert to orchestrate effective host defense. Whether and how these niche-derived immune cells participate in antiviral defense, particularly in determining infection outcomes, remains poorly understood.

To elucidate the mechanisms that determine life-or-death outcomes following viral infection, we took advantage of a binary outcome model in which virus-infected mice were categorized into recovering and non-recovering groups based on body weight loss. This approach enabled us to explore immune determinants that govern survival under apparently uniform genetic and environmental conditions. Comparative analyses revealed an enrichment of CXCR2^+^ cells resembling immunosuppressive myeloid-derived suppressor cells (MDSCs) at the initial site of infection in non-recovering mice. These cells exhibited immunosuppressive properties and were associated with attenuated inflammatory responses. Our findings suggest that these immunosuppressive CXCR2^+^ MDSC-like cells, likely derived from a local immune niche, may tip the balance between protective and detrimental immunity, ultimately determining survival or death following viral infection, pointing to an underappreciated layer of immune regulation.

## Results

### Suppressed inflammatory responses and enhanced viral dissemination underlie severe disease outcome in non-recovering mice

To explore determinants of disease severity independent of genetic or environmental factors, we first intranasally challenged mice with VSV and categorized them into two groups. Mice that retained ≥97% of their body weight from Day 5 to Day 6 were classified as “recovering mice” (R mice) with mild outcomes, while all other mice were classified as “non-recovering” mice (NR mice) with severe outcomes (Fig. 1a,b and Supplementary Fig. 1d,e). A detailed definition of these two groups is provided in the Methods.

**Fig 1.**
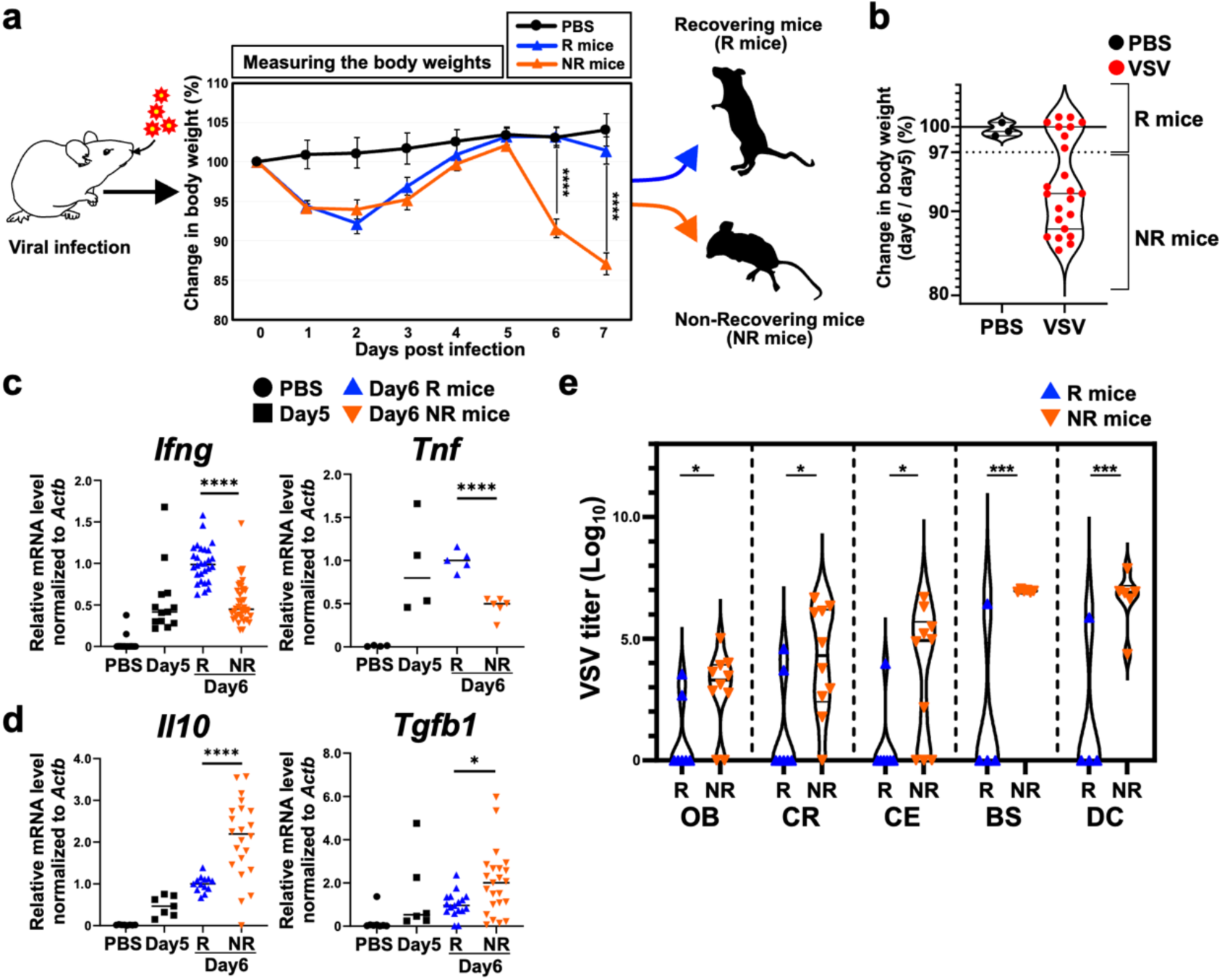
Non-recovering (NR) mice show an anti-inflammatory state. (**a**) General scheme showing the definition of recovering (R) mice and non-recovering (NR) mice after infection with VSV via the intranasal route. Change in body weight of uninfected mice (*n* = 3 mice), R mice (*n* = 8 mice), and NR mice (*n* = 15 mice). (**b**) Scatter plot depicting % body weight at 6 dpi compared with that at 5 dpi. Red dots indicate infected mice (*n* = 23 mice), and black dots indicate control mice (*n* = 3 mice). We categorized infected mice whose body weight at 6 dpi was ≥97% of that at 5 dpi as R mice, and others as NR mice. (**c,d**) mRNA levels of *Ifng*, *Tnf*, *Il10*, and *Tgfb1* in the olfactory bulb (OB) of uninfected mice (PBS), VSV-infected mice at 5 dpi (Day 5), R mice, and NR mice; *Ifng*: *n* = 16, 13, 28, and 38 mice, respectively; *Tnf*: *n* = 4, 4, 5, and 6 mice, respectively; *Il10*: *n* = 8, 7, 13, and 12 mice, respectively; and *Tgfb1*: *n* = 8, 6, 17, and 13 mice, respectively. (**e**) Viral titer in the OB, cortex (CR), cerebellum (CE), brainstem (BS), and diencephalon (DC) of R mice and NR mice at 6 dpi; OB, CR, and CE: *n* = 7 and 10 mice, respectively; BS an DC: *n* = 4 and 6 mice, respectively. Data are presented as mean ± SEM (a) or mean (c,d). *p < 0.05, ***p < 0.005, ****p < 0.001. Statistical analyses were performed using one-way ANOVA with Tukey’s post-tests (a,c,d) or unpaired two-sided Student’s t-tests (e). See also Supplementary Fig. 1.

Analysis of the olfactory bulb (OB), the initial infection site within the brain, revealed that levels of inflammatory cytokines such as *Ifng* and *Tnf*, which play pivotal roles in antiviral responses^10,22,23^, were significantly higher in R mice than in NR mice at 6 days post-infection (dpi) (Fig. 1c). By contrast, the levels of anti-inflammatory cytokines such as *Il10* and *Tgfb1*, which suppress antiviral immune responses and often protect against severe pathology^24,25^, were markedly lower in R mice than in NR mice (Fig. 1d). These results indicate that inflammatory responses to VSV infection are suppressed in NR mice. Consistent with these results, the viral titers in the brain were significantly higher in NR mice than in R mice (Fig. 1e). Notably, NR mice showed viral dissemination to distal brain regions such as the brainstem, which is a crucial regulator of respiratory and cardiovascular systems and is essential for survival^26^ (Fig. 1e). We detected no increase in cytokine expression, or VSV titers, in the lungs or liver from either group at 6 dpi (Supplementary Fig. 1f-h), indicating that the viral dissemination was limited to the brain at this phase of the infection. Taken together, these results suggest that “impaired immune responses and consequent enhanced viral spread” rather than “excessive immune responses” are associated with severe outcomes in this mouse model of VSV infection.

### CXCR2⁺ MDSC-like cells are more abundant in the OB of NR Mice

Next, we examined the mechanism underlying “impaired immune responses” in the OB of NR mice after the infection. We focused on differences in the profile of immune cells between R and NR mice by performing single-cell RNA sequencing (scRNA-seq) of CD45^+^ immune cells isolated from the OB at 6 dpi (Fig. 2a and Supplementary Fig. 2a,b), and found that CXCR2^+^ myeloid cells were more abundant in NR mice than in R mice (Fig. 2a,b and Supplementary Fig. 2c). We also found that although uninfected mice had very few CXCR2^+^ myeloid cells in the OB, their numbers increased markedly after infection with VSV (Fig. 2b). Importantly, these CXCR2^+^ myeloid cells expressed high levels of immunosuppressive molecules such as PD-L1 (*Cd274*) and *Acod1*, signature genes of immunosuppressive myeloid-derived suppressor cells (MDSCs)^27–30^ (Supplementary Fig. 2d). We thus termed them CXCR2^+^ MDSC-like cells in our study. In addition, we also observed that expression of *Cxcl1* and *Cxcl2*, both of which are ligands of CXCR2 (which is essential for infiltration of MDSCs into the tumor microenvironment^31,32^), increased in the OB after VSV infection, and was significantly higher in NR mice than in R mice (Fig. 2c). These findings suggest that CXCR2^+^ MDSC-like cells represent a population newly recruited to the infected site (particularly the OB) from external sources in response to viral infection, and that their pronounced accumulation in NR mice is associated with impaired immune responses.

**Fig 2.**
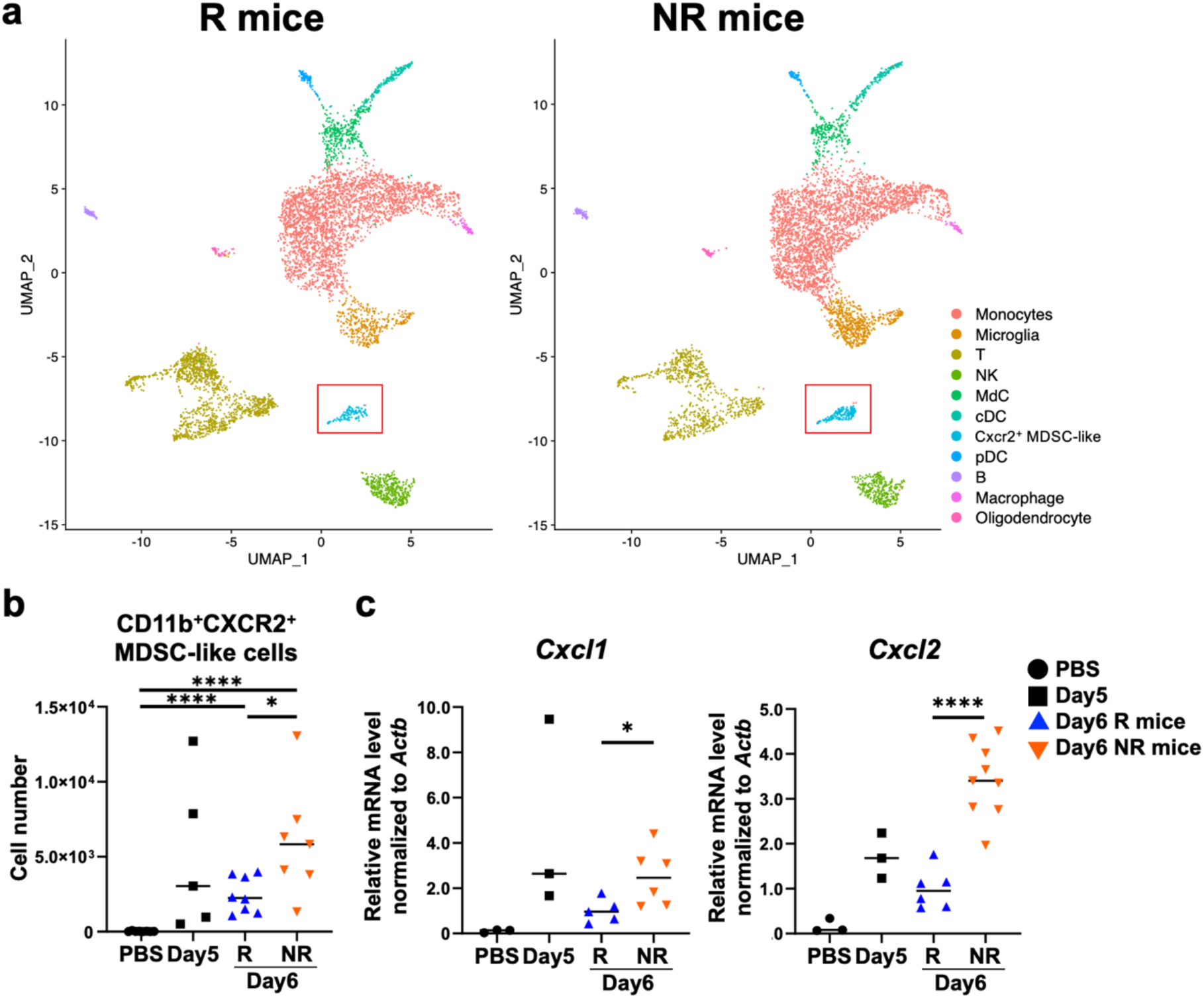
NR mice have more CXCR2^+^ MDSC-like cells. (**a**) UMAP of CD45^+^ immune cells in the OB from R mice (*n* = 3 mice) and NR mice (*n* = 3 mice) (cell numbers = 5,772 and 5,498 cells, respectively) at 6 dpi. pDC, plasmacytoid dendritic cell; cDC, conventional dendritic cell; MdC, monocyte derived cell. (**b**) Absolute number of CXCR2^+^ MDSC-like cells in the OB of uninfected mice (PBS) (*n* = 8 mice), VSV-infected mice at 5 dpi (Day 5) (*n* = 5 mice), R mice (*n* = 8 mice), and NR mice (*n* = 7 mice). (**c**) mRNA levels of *Cxcl1* and *Cxcl2* in the OB of PBS control, Day5, R mice, and NR mice; *Cxcl1*: *n* = 3, 3, 5, and 6 mice, respectively; *Cxcl2*: *n* = 3, 3, 6, and 9 mice, respectively. Data are presented as mean (b,c). *p < 0.05, ****p < 0.001. Statistical analyses were performed using one-way ANOVA with Tukey’s post-tests (b,c). See also Supplementary Fig. 2.

### CXCR2⁺ MDSC-like cells exhibit immunosuppressive activity upon viral infection

The apparent association between impaired immune responses and higher abundance of CXCR2^+^ MDSC-like cells in NR mice raises the possibility that these cells play a role in suppressing antiviral responses, thereby contributing to severe outcomes. Therefore, we utilized CXCR2 conditional knockout (cKO) mice (*Cxcr2*^fl/fl^; *Lysm*-Cre), which lack *Cxcr2* expression in myeloid cells, to inhibit the recruitment of CXCR2^+^ MDSC-like cells to the infected sites. At 6 dpi, the number of CXCR2^+^ MDSC-like cells, as well as expression of *Cxcr2*, were significantly lower in the OB of cKO mice than in that of control mice (*Cxcr2*^fl/fl^) (Fig. 3a,b). Furthermore, cKO of *Cxcr2* decreased expression of *S100a8* and *S100a9*, both of which were expressed exclusively by CXCR2^+^ MDSC-like cells among immune cells (Fig. 3b and Supplementary Fig. 2d), confirming that CXCR2^+^ MDSC-like cells infiltrate into the OB in a CXCR2-dependent manner after VSV infection. Viral titers in CXCR2 cKO mice were significantly lower than those in control mice, especially in the cortex, cerebellum, and diencephalon (Fig. 3c). Remarkably, expression levels of inflammatory cytokines in the OB of CXCR2 cKO mice, as well as survival rates, after VSV infection were significantly higher than those in control mice (Fig. 3d,f). Collectively, these findings suggest that CXCR2^+^ MDSC-like cells suppress inflammatory responses against the virus, leading to deleterious outcomes in NR mice.

**Fig 3.**
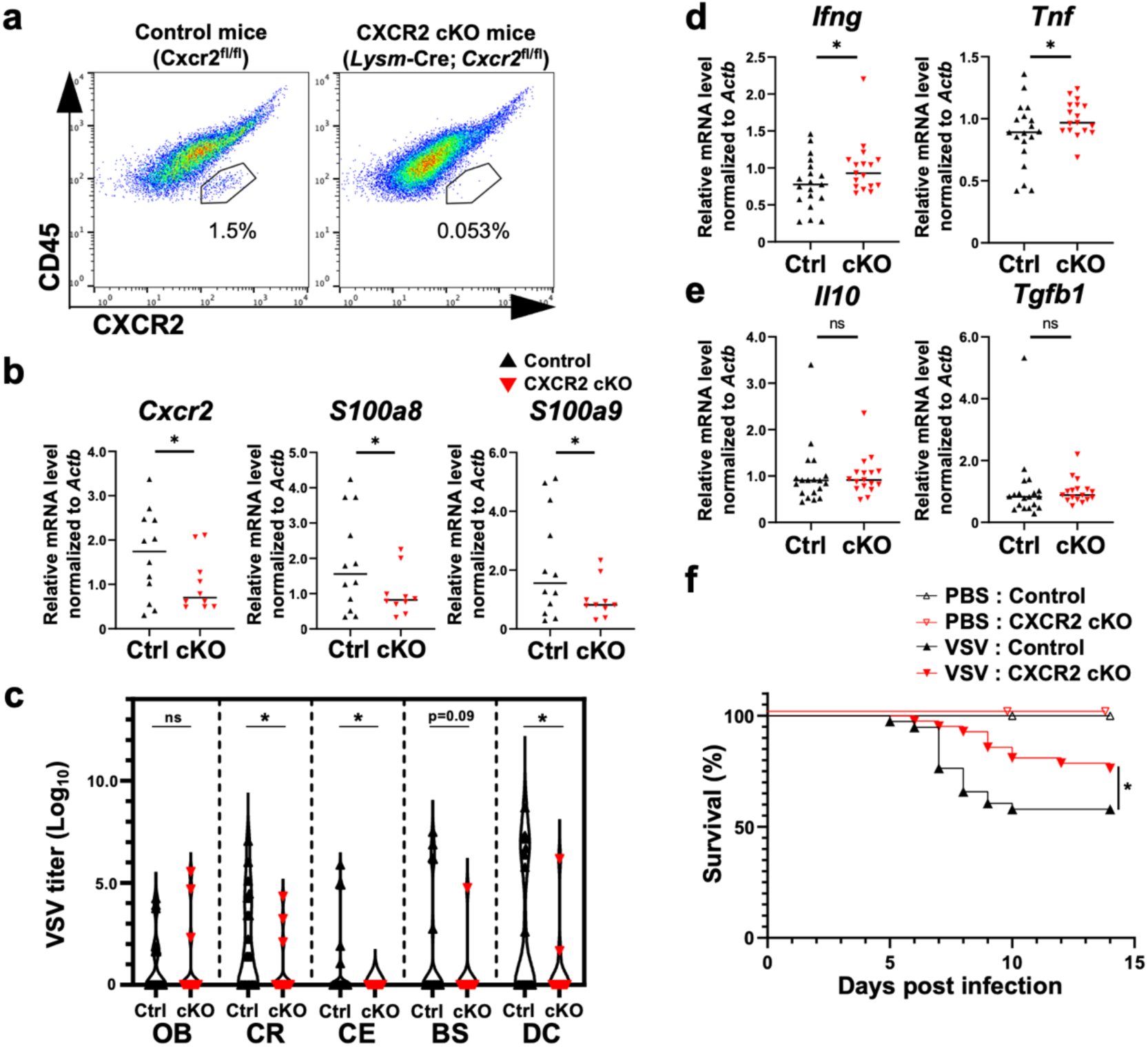
CXCR2^+^ MDSC-like cells suppress host defense. (**a**) Representative flow cytometry plots showing the percentage of CXCR2^+^ MDSC-like cells among CD45^+^ CD11b^+^ myeloid cells in the OB of control mice (*Cxcr2*^fl/fl^) and CXCR2 conditional knockout (cKO) mice (*Lysm*-Cre; *Cxcr2*^fl/fl^). (**b**) mRNA levels of *Cxcr2*, *S100a8*, and *S100a9* in the OB of control mice (*n* = 12 mice) and CXCR2 cKO mice (*n* = 10 mice). (**c**) Viral titer in each brain region of control mice (*n* = 20 mice) and CXCR2 cKO mice (*n* = 13 mice) at 6 dpi. (**d,e**) mRNA levels of *Ifng*, *Tnf*, *Il10*, and *Tgfb1* in the OB of control mice (*n* = 19 mice) and CXCR2 cKO mice (*n* = 17 mice). (**f**) Survival curve for uninfected (PBS) control mice (*n* = 9 mice), PBS CXCR2 cKO mice (*n* = 15 mice), VSV-infected control mice (*n* = 38 mice), and VSV-infected CXCR2 cKO mice (*n* = 42 mice). Data are presented as mean (b,d,e). ns, not significant, *p < 0.05, ****p < 0.001. Statistical analyses were performed using unpaired one-sided Student’s t-tests (b,c,d,e) or Log-rank test (f).

### CXCR2^+^ MDSC-like cells migrate to the OB from local niches

Next, we investigated the source of CXCR2^+^ MDSC-like cells. We parabiotically joined the circulation of wild-type (WT) mice to that of mice expressing the green fluorescent protein, kikGR, under the CAG promoter, thereby enabling us to distinguish recipient-derived and fluorescent donor-derived cells (Fig. 4a). While approximately 90% of microglia (resident immune cells in the brain) originated from recipient mice, about 45% of CD3e^+^ T cells, which are known to infiltrate from circulating blood following VSV infection, in the OB were derived from the donor mice (Fig. 4b), confirming the success of the surgery and shared circulation. Interestingly, only about 10% of CXCR2^+^ MDSC-like cells in the OB showed kikGR fluorescence after viral infection, which was comparable to those of microglia in the OB after viral infection (Fig. 4b), raising the possibility that these CXCR2^+^ MDSC-like cells originate from local niches in addition to the blood. Furthermore, analysis of differentially expressed genes and gene ontology pathways revealed upregulation of genes associated with “negative regulation of immune system processes” in CXCR2^+^ cells within the OB compared with CXCR2^+^ cells in the blood (Supplementary Fig. 3a,b). Downregulated pathways in CXCR2^+^ cells within the OB were largely related to chemotaxis, migration, and response to mechanical stimuli (Supplementary Fig. 3a,b). Taken together, these data indicate that CXCR2^+^ MDSC-like cells in the OB have a role largely distinct from that of CXCR2^+^ cells in the blood following viral infection.

**Fig 4.**
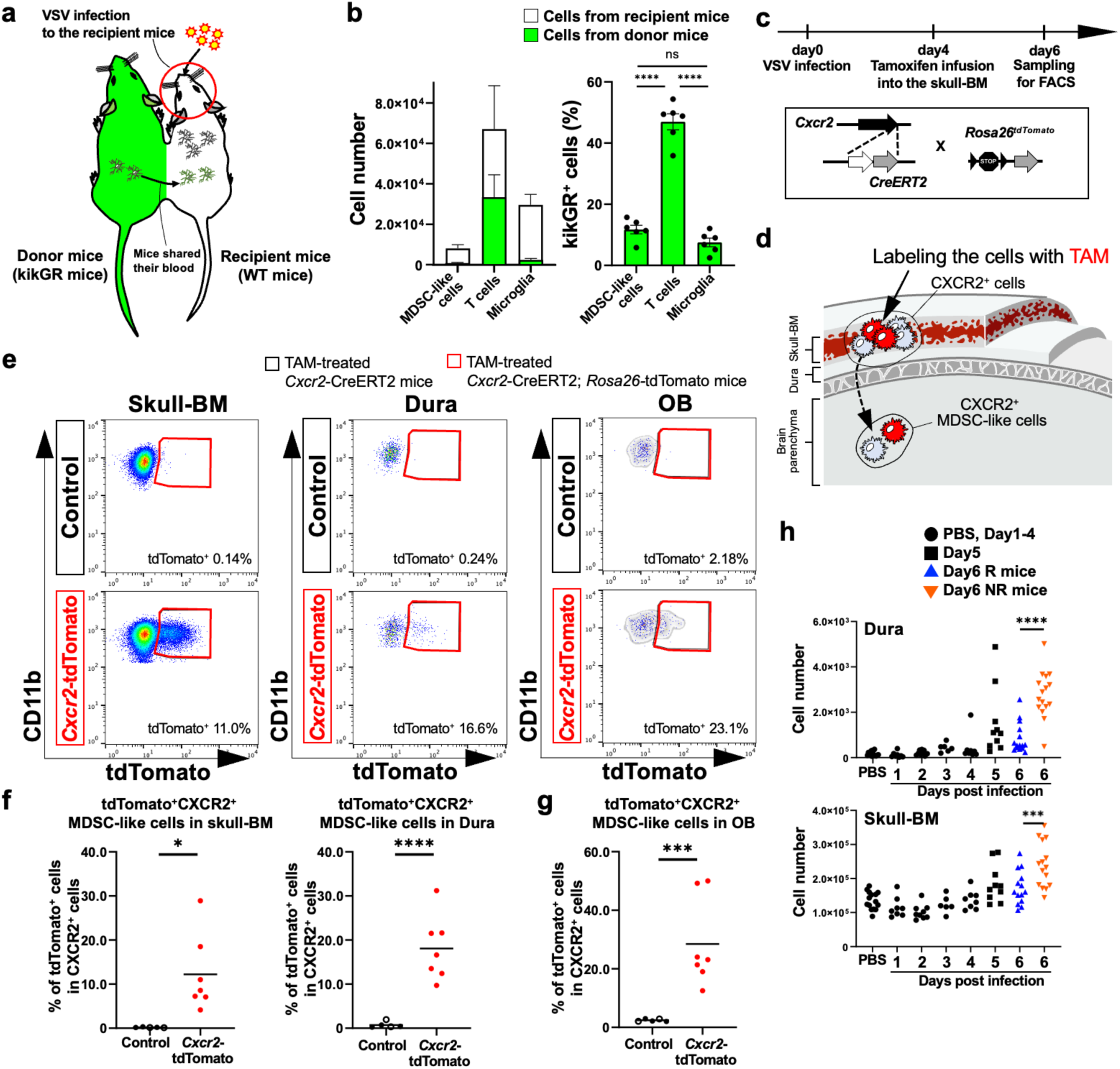
CXCR2^+^ MDSC-like cells migrate to the OB from local niche. (**a**) General scheme of parabiosis experiment. (**b**) Absolute cell number and percentage of kikGR^+^ cells among CXCR2^+^ MDSC-like cells, CD3e^+^ T cells, and CD11b^+^ Ly6c^-^ microglia in the OB at 6 dpi (*n* = 6 mice). (**c,d**) General scheme showing the method used to label Cxcr2^+^ cells in the skull-BM with tdTomato. Tamoxifen (TAM) (20 mg/ml in 30 μl of corn oil) was placed on the surface of the skull at 4 dpi. (**e**) Representative flow cytometry plots showing the percentage of tdTomato^+^ cells among CD45^+^ CD11b^+^ CXCR2^+^ myeloid cells in the OB, dura, and skull-BM of TAM-treated *Cxcr2*-CreERT2 (Control) mice (*n* = 3 mice) and TAM-treated *Cxcr2*-CreERT2; *Rosa*-tdTomato (*Cxcr2*-tdTomato) mice (*n* = 14 mice). (**f,g**) Percentage of tdTomato^+^ cells among CD45^+^ CD11b^+^ CXCR2^+^ myeloid cells in the OB, dura, and skull-BM of Control mice (*n* = 3 mice) and *Cxcr2*-tdTomato mice (*n* = 14 mice). (**h**) Absolute number of CXCR2^+^ cells in the dura and skull-BM from uninfected mice (PBS; *n* = 11 mice), 1 dpi (Day 1; *n* = 12 mice), 2 dpi (Day 2; *n* = 9 mice), 3 dpi (Day 3; *n* = 6 mice), 4 dpi (Day 4; *n* = 8 mice), 5 dpi (Day 5; *n* = 12 mice), R mice (*n* = 15 mice), and NR mice (*n* = 14 mice). Data are presented as mean ± SEM (b) or mean (f,g). *p < 0.05, **p < 0.01, ***p < 0.005, ****p < 0.001. Statistical analyses were performed using one-way ANOVA with Tukey’s post-tests (b,h) or paired one-sided t-tests (f,g). See also Supplementary Fig. 3 and Supplementary Fig. 4.

We then further explored the possibility that MDSC-like cells originate from local niches. Recent studies identified the skull-bone marrow (skull-BM) as a unique local source of immune cells that migrate directly to the brain parenchyma through the dura^21,33,34^. To determine whether CXCR2^+^ MDSC-like cells infiltrate into the OB from the skull-BM after VSV infection, we locally treated tamoxifen (TAM) onto the skull surface of *Cxcr2*-tdTomato mice (*Cxcr2*-CreERT2; *Rosa26*-IRES-tdTomato) and successfully labeled CXCR2^+^ cells in the skull-BM (Fig. 4c-f). Remarkably, at 6 dpi, we observed tdTomato^+^ cells among CXCR2^+^ cells in the dura and the OB of *Cxcr2*-tdTomato mice, whereas tdTomato^+^ cells were scarcely detected in the OB of either control mice (*Cxcr2*-CreERT2) or uninfected *Cxcr2*-tdTomato mice (Fig. 4e-g and Supplementary Fig. 4a). These results indicated that skull-BM-derived CXCR2⁺ cells migrate into the brain parenchyma upon viral infection. Interestingly, during the viral infection, the numbers of CXCR2^+^ cells in both the skull-BM and dura increased gradually from 4 dpi, and were significantly higher in NR mice than in R mice at 6 dpi (Fig. 4h). Together, these findings suggest that a significant fraction of CXCR2⁺ MDSC-like cells originates from the local niche of the skull-BM, and that enhanced production of these cells in the skull-BM of NR mice is associated with more severe disease outcomes following viral infection.

### Intracisternal administration of a CXCR2 inhibitor improves outcomes after viral infection

Finally, we investigated whether skull-BM-derived CXCR2^+^ MDSC-like cells suppress antiviral responses. To prevent migration of CXCR2^+^ MDSC-like cells from the skull-BM to the brain parenchyma, we administered the CXCR2 inhibitor (SB225002) to mice via the cisterna magna after VSV infection (Fig. 5a). We found that intracisternal magna (ICM) administration of the CXCR2 inhibitor significantly reduced the number of CXCR2^+^ MDSC-like cells in the OB compared with control treatment (Fig. 5b). Importantly, ICM administration of the CXCR2 inhibitor increased expression of inflammatory cytokines and decreased that of anti-inflammatory cytokines in the OB after VSV infection, compared with the control treatment (Fig. 5c,d). Furthermore, the CXCR2 inhibitor treatment significantly improved survival rates after VSV infection, with all inhibitor-treated mice surviving throughout the observation period (Fig. 5e). By contrast, intraperitoneal (IP) administration of the same CXCR2 inhibitor, which blocks migration of CXCR2^+^ MDSC-like cells from the circulating blood, did not improve survival rates after VSV infection (Supplementary Fig. 5a,b). Taken together, these findings suggest that CXCR2^+^ MDSC-like cells derived from the skull-BM play a critical role in suppressing host defenses against the virus, ultimately reducing survival.

**Fig 5.**
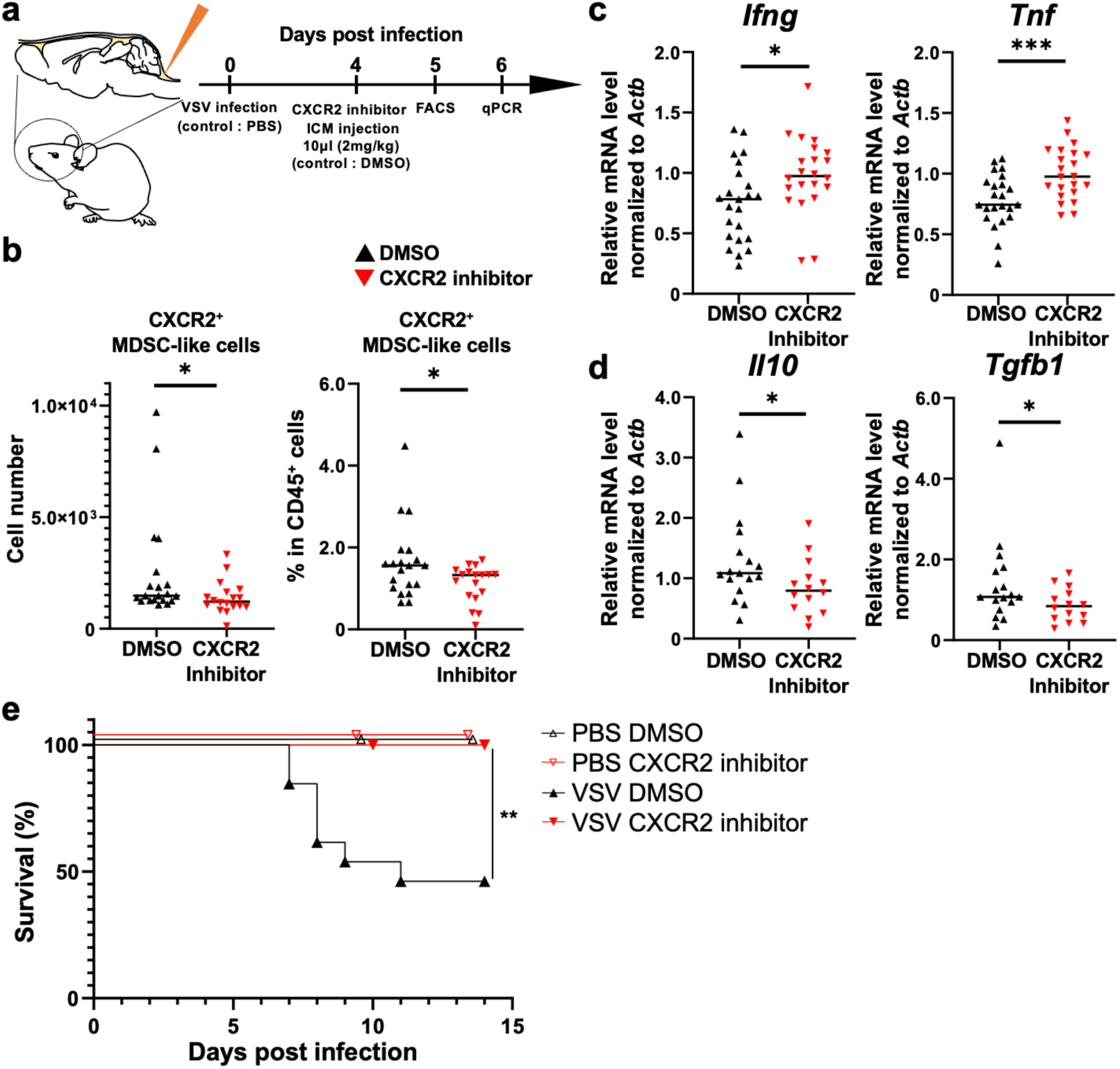
Intracisternal administration of a CXCR2 inhibitor improves outcomes after viral infection. (**a**) General scheme of the CXCR2 inhibitor injection. ICM, intracisterna magna; DMSO, dimethyl sulfoxide. (**b**) Absolute number and percentage of CXCR2^+^ MDSC-like cells among CD45^+^ immune cells in control (DMSO) mice (*n* = 20 mice) and CXCR2 inhibitor-treated (CXCR2 inhibitor) mice (*n* = 18 mice) at 5 dpi. (**c,d**) mRNA levels of *Ifng*, *Tnf*, *Il10*, and *Tgfb1* in the OB of DMSO mice and CXCR2 inhibitor mice at 6 dpi; *Ifng* and *Tnf*: *n* = 22 and 24 mice, respectively; *Il10* and *Tgfb1*: *n* = 16 and 16 mice, respectively. (**e**) Survival curves for PBS DMSO mice (*n* = 3 mice), PBS CXCR2 inhibitor mice (*n* = 3 mice), VSV-infected DMSO mice (*n* = 14 mice), and VSV-infected CXCR2 inhibitor mice (*n* = 12 mice) after ICM injection. Data are presented as mean (b,c,d). *p < 0.05, **p < 0.01, ***p < 0.005. Statistical analyses were performed using unpaired one-sided Student’s t-tests (b,c,d) or log-rank test (e). See also Supplementary Fig. 5.

## Discussion

Using a mouse model that eliminates differences in pre-existing conditions such as genetic background and environmental exposures, we found that, contrary to the prevailing focus on immunopathology driven by excessive immune activation, immune suppression—specifically mediated by CXCR2⁺ MDSC-like cells at the initial site of infection—determines lethality in this setting. These cells accumulated prominently in non-recovering animals, and their local, but not systemic, blockade of their CXCR2-dependent recruitment improved survival. These findings indicate that immunosuppressive activity by CXCR2⁺ MDSC-like cells, likely originating from a local immune niche, plays a critical role in shaping infection outcomes. Notably, this immunoregulatory function appears to be distinct from that of FOXP3⁺ Tregs, which are known to contribute to immune suppression in the brain^35–37^, as these cells neither expanded nor altered inflammation when depleted upon viral infection (Supplementary Fig. 6). These results underscore the functional predominance of CXCR2⁺ MDSC-like cells in this context.

Beyond their immunosuppressive function, CXCR2⁺ MDSC-like cells exhibited features that distinguish them from conventional MDSCs. Specifically, while they expressed anti-inflammatory cytokines such as *Il10* and *Tgfb1*, they lacked canonical markers typically associated with conventional MDSCs, including *Ly6c* and *Ly6g*^38^. Furthermore, unlike tumor-associated MDSCs that are commonly recruited from the circulation^39^, a significant fraction of CXCR2⁺ MDSC-like cells in our study originated from the skull bone marrow—a local hematopoietic niche directly connected to the central nervous system. This distinct anatomical origin, together with a partially overlapping yet divergent gene expression profile, suggests that the CXCR2⁺ MDSC-like cells described here constitute a previously unrecognized population. These findings provide a new conceptual framework in which local immune niches serve as reservoirs for immunosuppressive cells that shape infection outcomes.

The CXCR2⁺ MDSC-like cells appear to expand within a local niche in response to brain infection and are likely mobilized by elevated levels of CXCL1 and CXCL2. However, the route by which they are recruited remains unclear. Notably, recent studies have shown that cerebrospinal fluid (CSF) can access the skull bone marrow through direct channels traversing the dura mater, thereby enabling CSF-borne signals to influence hematopoietic activity in the marrow^33,40,41^. It is plausible that CXCL1 and CXCL2 reach the skull marrow through these channels, triggering the mobilization of CXCR2⁺ MDSC-like cells. It will be important to determine whether CXCL1 and CXCL2 reach the skull bone marrow through dural channels to promote recruitment of these cells. Additionally, whether CSF contributes to their expansion—and what CSF-borne signals mediate this process—represents an important direction for future study.

Our findings demonstrate that local administration of a CXCR2 inhibitor enhances inflammation and improves disease outcome (Fig. 5e and Supplementary Fig. 7). This approach represents a novel form of host-targeted therapy (HTT)—one that promotes viral clearance by relieving immunosuppression within a localized immune niche. This HTT strategy is mechanistically distinct from conventional direct-acting antivirals (DAAs), which aim to inhibit viral replication directly, as well as from systemic immunostimulatory agents, which broadly activate host defenses. Therefore, CXCR2 inhibition may offer a complementary therapeutic avenue that enhances antiviral efficacy.

The immunosuppressive function of these CXCR2⁺ myeloid cells may have broader relevance beyond viral infection. While they may dampen protective immunity in the context of infection, they could conversely support resolution and recovery in chronic inflammatory conditions. Indeed, our analysis of human clinical trial data suggests that administration of the CXCR2 inhibitor navarixin—originally developed to block inflammatory neutrophils—may exacerbate inflammatory conditions such as nasopharyngitis in patients with asthma or chronic obstructive pulmonary disease (COPD) (Supplementary Fig. 8), potentially due to unintended inhibition of immunosuppressive CXCR2⁺ myeloid cells. These findings underscore the importance of further characterizing this population to guide the rational design of anti-inflammatory therapies.

## Acknowledgments

The authors thank Shohei Hori for helpful discussions, suggestions, *Foxp3*-GFP mice and *Foxp3*-DTR; Maki Ukai-Tadenuma and Hidemi Kasahara for generating the *Cxcr2*-CreERT2 mice; Kaoru Tabata for handling the mouse colony; Athushi Miyawaki, Junichi Miyazaki for kikGR mice; and all members of Laboratory of Molecular Biology (Graduate School of Pharmaceutical Sciences, The University of Tokyo) community for feedback throughout this project.

## Author contributions

A.S. inputted conceptualization, designed the methods, and performed analysis. A.S. and R.S. acquired samples. H.S. prepared the platforms of RNA-seq and proteome analysis. T.M. analyzed the clinical trial data. S.S. and K.M. helped with scRNA-seq analysis. H.U. generated the transgenic mouse line *Cxcr2*-CreERT2. M.Y. generated transgenic mouse line *Rosa26*-CS-lsl-DTR-IRES-tdTomato mice. Y.G. and T.O. supervised all work. A.S., R.S., Y.G., and T.O. wrote the paper with input from all authors. All co-authors read, reviewed, and approved the manuscript.

## Data availability

The source data are provided within the paper. The codes used for analyses of published mouse OB transcriptome data are available at https://github.com/satomindog/MDSC-like.

## Additional information

Supplemental information is available for this paper.

## Methods

### Mice

C57BL/6 were purchased from CLEA Japan. CAG-kikGR mice (B6.Cg-Gt(ROSA)26Sor<tm1.1(CAG-kikGR)Kgwa>) was provided by the RIKEN BRC through the National Bio Resource Project of the MEXT, Japan^45–47^. *Foxp3*-GFP mice^48^ and *Foxp3*-DTR mice (JAX 016958) were obtained from S. Hori (Graduate School of Pharmaceutical Sciences, The University of Tokyo, Japan). Treg ablation in *Foxp3*-DTR mice was induced by intraperitoneal (IP) injection of diphtheria toxin (DT) (diluted in PBS, 25 ug/kg) every other day from one day before VSV infection. Transgenic mouse line *Cxcr2*^fl/fl^ homozygous mice (JAX 024638) and *Lysm*-Cre heterozygous mice (JAX 004781) were purchased from the Jackson Laboratory and mated to produce control and knockout littermates. Transgenic mouse line *Cxcr2*-CreERT2 (Accession No. IRCN-GE0057: https://core.ircn.jp/en/es-virus-core/) was generated as described previously^49^, and as described below. *Rosa26*-CS-lsl-DTR-IRES-tdTomato mice^43^ were provided by M. Yanagisawa (International Institute for Integrative Sleep Medicine (WPI-IIIS), The University of Tsukuba). *Cxcr2*-CreERT2 mice and *Rosa26*-CS-lsl-DTR-IRES-tdTomato mice were mated and the littermates used for labeling CXCR2^+^ cells in the skull-BM via tamoxifen-treatment. All mice were maintained in a temperature- and humidity-controlled environment (23°C ± 3°C and 50 ± 15%) under a 12-h-light/12-h-dark cycle.

Mice were housed up to six mice per cage (Innocage, Innovive or Micro BARRIER Systems, Edstrom Japan) containing chips (PALSOFT, Oriental Yeast), and with access to irradiated food (CE-2, CLEA Japan) and water *ad libitum*. Eight-week-old female mice were used for all experiments. Intranasal infection with VSV (serotype New Jersey) at a titer of 1.0 × 10^7^ plaque-forming units in 50 µl of PBS per nostril was performed under anesthesia with 5% isoflurane. All animal procedures and experiments were approved by and performed in accordance with the guidelines of the Animal Care and Use Committee of The University of Tokyo.

### Generation of transgenic mouse lines

To construct the donor vector, a 4,000-bp DNA fragment upstream from the stop codon of *Cxcr2*, and a 3,969-bp downstream fragment that included the stop codon, were amplified from the genomic DNA of C57BL/6 mice and used as homologous arms. A mouse codon-optimized P2A self-cleavage sequence (19 amino acids) and CreERT2 were inserted in-frame with *Cxcr2* via a glycine-serine-glycine linker placed between the homologous arms. A woodchuck hepatitis virus post-transcriptional regulatory element and an SV40 late polyadenylation signal were added downstream of CreERT2.

Additionally, a 1.7-kb fragment containing an FRT-PGK promoter-puromycin resistance gene-polyA-FRT cassette was inserted after the SV40 polyadenylation signal for use as a selection marker. The circular donor vector was co-transfected embryonic stem (ES) cells derived from C57BL/6 mice along with TALEN expression vector pairs targeting 5’-agctcgtcttcagcaaaca-3’ and 5’-ttaggtgaacagtctt-3’ to establish *Cxcr2*-CreERT2 knock-in (KI) ES cells. Correctly targeted ES cell clones were injected into eight-cell-stage ICR mouse embryos, which were cultured to the blastocyst stage and transferred into pseudopregnant ICR females. The resulting mice, entirely derived from KI-ES cells, were used as *Cxcr2*-CreERT2 mice in the experiments.

### Definition of R mice and NR mice after viral infection

Some mice succumbed by 10 days post-infection (dpi) with VSV, while others survived beyond 10 dpi (Supplementary Fig. 1a). In this study, body weight was the parameter used to assess the degree of disease severity after viral infection. First, VSV-infected mice were categorized into two groups: mice that died by 10 dpi, and mice that survived beyond 10 dpi. Comparison of the body weight of deceased and surviving mice revealed that changes in the body weight between 5–8 dpi were significantly different between the two groups (Supplementary Fig. 1b,c). Since more than half of deceased mice had already died by 7 dpi (Supplementary Fig. 1a), changes in body weight during 5–6 dpi were used to classify VSV-infected mice into two new groups: recovering mice (R mice) and non-recovering mice (NR mice). Changes in body weight were calculated as follows:

X (%)= {1+ ((”The body weight at 6 dpi”-”The body weight at 5 dpi”))/(”The body weight at 5 dpi”)}× 100

Since an X value of 97% corresponded to the highest survival rate for R mice, and the lowest for NR mice, at 6 dpi, mice with X ≥ 97% were categorized as R mice, and those with X < 97% were categorized as NR mice (Fig. 1a,b and Supplementary Fig. 1d,e).

### Preparation and quantification of VSV

Virus stocks were grown and viral titers quantified as described previously^50^. For the in vivo studies, tissues were collected without perfusion, immersed in 500 μl of PBS, and homogenized in a multi-beads shocker (Yasui Kikai). Homogenates were transferred to a new 1.5 ml tube and spun for 5 min at 8,000 rpm at 4°C. Supernatants were filtered through a 0.45 μm filter to remove cell debris prior to quantification.

### Isolation and flow cytometry analysis of immune cells

After perfusion with 10 ml of PBS, the OB, dura, and skull-BM were collected from mice anesthetized with 150 µl of combination anesthetic (containing 0.75 µg/µl Medetomidine Hydrochloride, 0.4 µg/µl Midazolam, and 0.4 µg/µl Butorphanol Tartrate). The OB and calvarium were cut into small pieces using a knife or sterile scissors, and then suspended at 37°C for 15 min in 200 µl of Collagenase Ⅷ (1 mg/ml, containing 0.5 mg/ml DNaseⅠ and 2% FBS) in 1.5 ml low-cell binding tubes. After centrifugation for 5 min at 300 × g/4°C, the cells were resuspended in 10% Percoll (Sigma Aldrich) in 0.2% BSA/PBS and then filtered through a 35 µm mesh filter (Falcon) into a new 1.5 ml low-cell binding tube. After centrifugation for 5 min at 300 × g/4°C, the cells were incubated at 4°C for 15 min with Microbeads anti-CD45 (1:100, clone 30F11, Miltenyi Biotec, 120-008-885). After incubation and centrifugation, the samples were subjected to Magnetic Activated Cell Sorting (MACS). Cells trapped by MACS were centrifuged for 5 min at 2,500 rpm/4°C and then incubated at 4°C for 15 min with 4% paraformaldehyde (PFA). After centrifugation for 5 min at 800 × g/4°C, the cell suspensions were washed and then stained at 4°C for 30 min with the following anti-mouse antibodies: anti-CD45.2 PerCP/Cy5.5 (1:200, clone 104, BioLegend, 109828), anti-CD11b APC/Cyanine7 (1:500, clone M1/70, BioLegend, 101226), anti-CD3 Pacific Blue (1:200, clone 17A2, BioLegend, 100214), anti-CD16/32 (1:200, clone 93, BioLegend, 101302), anti-CD182(CXCR2) PE (1:200, clone SA044G4, BioLegend, 149304), anti-Ly6C APC (1:200, clone HK1.4, BioLegend, 128016), CD182(CXCR2) APC (1:200, clone SA044G4, BioLegend, 149311), and anti-CD25 APC (1:200, clone PC16, BioLegend, 102012). The samples were then washed and acquired immediately using a FACS Aria flow cytometer (Becton Dickinson). Data were analyzed using FlowJo software.

### Single-cell RNA sequencing and analysis

The OB was collected from R mice (n = 3) and NR mice (n = 3) at 6 dpi. CD45^+^ immune cells were isolated by MACS using an anti-CD45.2 PE/Cyanine7 antibody (1:100, clone 104, BioLegend, 109829) and a Microbeads anti-PE antibody (1:100, Miltenyi Biotec, 130-048-801). Single-cell RNA-seq (scRNA-seq) libraries were prepared in accordance with the BD Rhapsody protocol. Each sample was stained with the following antibodies: anti-mouse Hashtag 7 (1:200, clone M1/42, 30-F11, BioLegend, 155813), anti-mouse Hashtag 7 (1:200, clone M1/42, 30-F11, BioLegend, 155813), anti-mouse Hashtag 8 (1:200, clone M1/42, 30-F11, BioLegend, 155845), anti-mouse Hashtag 9 (1:200, clone M1/42, 30-F11, BioLegend, 155877), anti-mouse Hashtag 10 (1:200, clone M1/42, 30-F11, BioLegend, 155879), anti-mouse Hashtag 11 (1:200, clone M1/42, 30-F11, BioLegend, 155881), and anti-mouse Hashtag 12 (1:200, clone M1/42, 30-F11, BioLegend, 155823). The sequence was analyzed by ImmunoGeneTeqs (Chiba, Japan) using the TAS-Seq platform^51^. In total, 11,270 single cells were sequenced: 5,772 from R mice and 5,498 cells from NR mice. Cells expressing < 200 features, > 2,500 features, or > 5% mitochondrial genes were excluded from the analysis. Dimensionality reduction and clustering analysis were performed using Seurat v4.3.0. The following cells were annotated using conventional cell markers: monocytes (*Ly6c1*), microglia (*Hexb*, *Cx3cr1*), T cells (*Cd3e*), NK cells (*Klrb1c*), MdC (*Cd86*), cDC (*Ccr7*), MDSC-like (*Cxcr2*), pDC (*Siglech*), B cells (*Cd19*), macrophages (*Mrc1*, *Cx3cr1*), and oligodendrocytes (*Plp1*).

### RNA extraction, reverse transcription, and qPCR

Total RNA was isolated from the tissues of infected mice using the RNAiso Plus reagent (Takara) after perfusion with 10 ml of PBS to remove intravascular cells. RNA extraction and real-time PCR analysis were performed as previously described^52^ to measure gene expression. The amount of target mRNA was normalized to that of *Actb* RNA. The sense and antisense primers, respectively, were as follows: mouse *Actb*, 5’-AATAGTCATTCCAAGTATCCATGAAA-3’ and 5’-GCGACCATCCTCCTCTTAG-3’; mouse *Ifng*, 5’-AGCTCTGAGACAATGAACG-3’ and 5’-GTGGCAGTAACAGCCAGAA-3’; mouse *Tnf*, 5’-TGGCCTCCCTCTCATCAGTT-3’ and 5’-GCTTGTGACTCGAATTTTGAGAAG-3’; mouse *Il10*, 5’-TCCTTGATTTCTGGGCCAT-3’ and 5’-GCAGGACTTTAAGGGTTACT-3’; mouse *Tgfb1*, 5’-CATTAAGGAGTCGGTTAGCAG-3’ and 5’-AACTGCACCCACTTCCCAGTC-3’; mouse *Foxp3*, 5’-TGCCTCCTCCAGAGAGAAGTG-3’ and 5’-TTGGCCAGCGCCATCTT-3’.

### Parabiosis surgery

Mice were anesthetized with 150 µl of a combination anesthetic (containing 0.75 µg/µl Medetomidine Hydrochloride, 0.4 µg/µl Midazolam, and 0.4 µg/µl Butorphanol Tartrate) prior to parabiosis surgery. Parabiosis between an 8-week-old WT female mouse and a CAG-kikGR female mouse was performed as described previously^53^. Briefly, the skin from the leg joint to the arm joint was cut, and each joint from one mouse was tied to that of the other using surgical sutures. The skin was then sutured together to yield parabiotic mice. These mice we maintained for 1 month prior to viral infection. Consistent with a previous finding that half of CD45^+^ CD11b^-^ lymphoid cells in the spleen are replaced by donor-derived cells by 14 days post-parabiosis^54^, we observed that approximately half of CD45^+^ CD11b^-^ lymphoid cells in the spleen of WT mice were donor-derived and showing green fluorescence, kikGR, (data not shown), confirming successful connection of the bloodstreams of the recipient and donor mice.

### Intracisterna magna (ICM) injection

Mice were anesthetized with 5% isoflurane, placed on stereotactic frames, and the CXCR2 inhibitor was injected into the cisterna magna. At 4 dpi, 10 µl of CXCR2 inhibitor (2 mg/kg; SB225002) in artificial cerebrospinal fluid (aCSF, containing 5% Tween, 4% Fast Green, and 20% polyethylene glycol) was administrated at a rate of 2 µl/min using glass capillary. Following injection, the needle was left in place for 2 min to avoid backflow. Then, the mice were sutured, placed on a heating pad, and allowed to recover for 1 h.

### Labeling of skull-BM

Mice were anesthetized with 5% isoflurane and placed on stereotactic frames. The skull surface was exposed and cleaned with 70% ethanol. To label cells in the skull-BM using lentivirus, the lentivirus was suspended in 20 µl of aCSF reagent and placed onto the skull surface, centered on the bregma, for 15 min. The same procedure was used to label labeling CXCR2^+^ cells in the skull-BM with 20 mg/ml tamoxifen in corn oil. After the skin was sutured, the mice were placed on a heat pad and allowed to recover for 1 h.

### RNA sequencing (FLASH-seq) analysis

During the flow cytometry procedure, dithio-bis (succinimidyl propioneta; DSP) was filtered through a 20 µm mesh filter (pluriSelect) and used instead of 4% PFA for cell fixation prior to extraction of RNA. The cells were incubated with DSP at room temperature for 30 min. Next, 2,000 CD45^+^ CD11b^+^ CXCR2^+^ cells were sorted by FACS and collected into 3.82 µl of lysis buffer. Template preparation was conducted using FLASH-seq as described previously^55,56^. Sequencing was carried out on an Illumina NextSeq 2000 platform with paired-end 36 bp reads. Sequence reads aligned to the mouse reference genome (mm10) using Hisat2 (v2.2.1). Row read counts of each gene was calculated using featureCounts (v2.0.1) and were normalized as Transcripts Per Kilobase Million (TPM). Gene ontology analysis was performed using Metascape. Dot plots and volcano plots were generated using ggplot within the ggplot2 package 3.3.6.

### Investigation of the inflammatory effects of navarixin in clinical trials

First, the result_group and reported_event tables were obtained from AACT (https://aact.ctti-clinicaltrials.org/) and integrated. Records containing “navarixin” in the title were selected, and the corresponding clinical trial IDs (nct_id_x) were retrieved. From the adverse_event_term entries linked to the filtered nct_id_x, those categorized under organ_system as “Infections and infestations” were extracted. Among these terms, inflammation-related entries, namely “Nasopharyngitis,” “Rhinitis,” “Diverticulitis,” “Otitis media chronic,” and “Gastroenteritis”, were chosen. In addition to navarixin, albuterol and formoterol were used as control drugs due to their similarity in clinical usage. For comparison, the adverse event terms were restricted to nasopharyngitis. Finally, 41 records from 25 clinical studies were obtained. To elucidate the inflammatory effects of navarixin, the following analyses of curated data were conducted using a nested bootstrap approach: first, data were separated by term, then for each clinical trial, the risk ratio was calculated through bootstrapping within the trial, and the results were integrated. Risk ratios across trials were then combined via bootstrapping, and 95% confidence intervals were calculated. The number of bootstrap iterations was set to 5,000. All analyses were performed using Python 3.10.

### Statistical analysis and reproducibility

Statistical analysis was performed using GraphPad Prism Software. Data from more than three independent groups were analyzed by one-way analysis of variance (ANOVA) with Tukey’s post-tests. For the volcano plot depicting -fold changes in body weight, FDR-adjusted Wald’s test was used (R package DESeq2). Two-group comparisons were made using two-tailed unpaired Student’s t-tests, one-tailed unpaired Student’s t-tests, or one-sided paired Student’s t-tests. For pooled analysis of results from independent repeats, all mice from the same experimental group were normalized to the average levels for R mice or control mice, and a new statistical comparison was made for the entire pooled experiment. *P < 0.05, **P < 0.01, ***P < 0.005,****P < 0.001, and ns; not significant. All experiments were repeated between two and eight times.

## Supplementary Figures and Legends

**Supplementary Fig 1.**
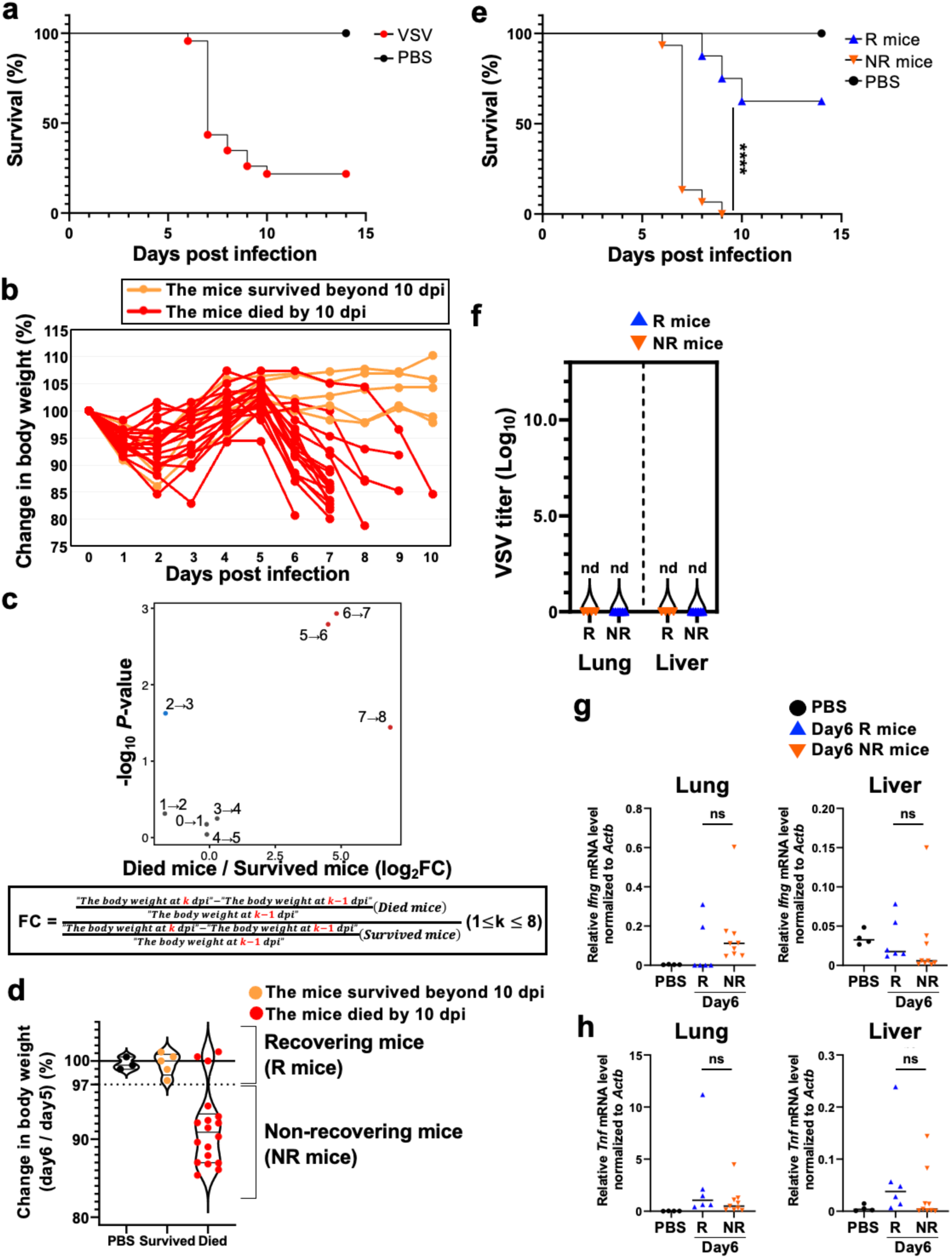
VSV-infected mice are categorized into R mice and NR mice, related to Fig. 1. (**a**) Survival curves for uninfected (PBS) mice (*n* = 3 mice) and VSV-infected (VSV) mice (*n* = 23 mice). (**b**) Changes in the body weight of VSV-infected mice that survived beyond 10 dpi (orange, *n* = 5 mice), and of those that died by 10 dpi (red, *n* = 18 mice). (**c**) The fold change (FC) in body weight was defined as shown. A lof2FC>|1| and an FDR-adjusted Wald’s test of P<0.05 are colored red and blue, respectively. The red dots indicate lof2FC>1, and the blue dots indicate lof2FC<-1. (**d**) Scatter plot depicting % body weight at 6 dpi compared with that at 5 dpi for uninfected mice (*n* = 3 mice), VSV-infected mice that survived beyond 10 dpi (orange, *n* = 5 mice), and those that died by 10 dpi (red, *n* = 18 mice). (**e**) Survival curves for uninfected (PBS) mice (*n* = 3 mice), recovering (R) mice (*n* = 8 mice), and non-recovering (NR) mice (*n* = 15 mice). (**f**) Viral titers in the lungs and liver of R mice (*n* = 4 mice) and NR mice (*n* = 6 mice) at 6 dpi. (**g,h**) mRNA levels of *Ifng* and *Tnf* in the lung or liver of PBS mice (*n* = 4 mice), R mice (*n* = 6 mice), and NR mice (*n* = 9 mice). Data are presented as mean (g,h). nd, no data, ns, not significant, ****p < 0.001. Statistical analyses were performed using log-rank test (e) or unpaired two-sided Student’s t-tests (g,h).

**Supplementary Fig 2.**
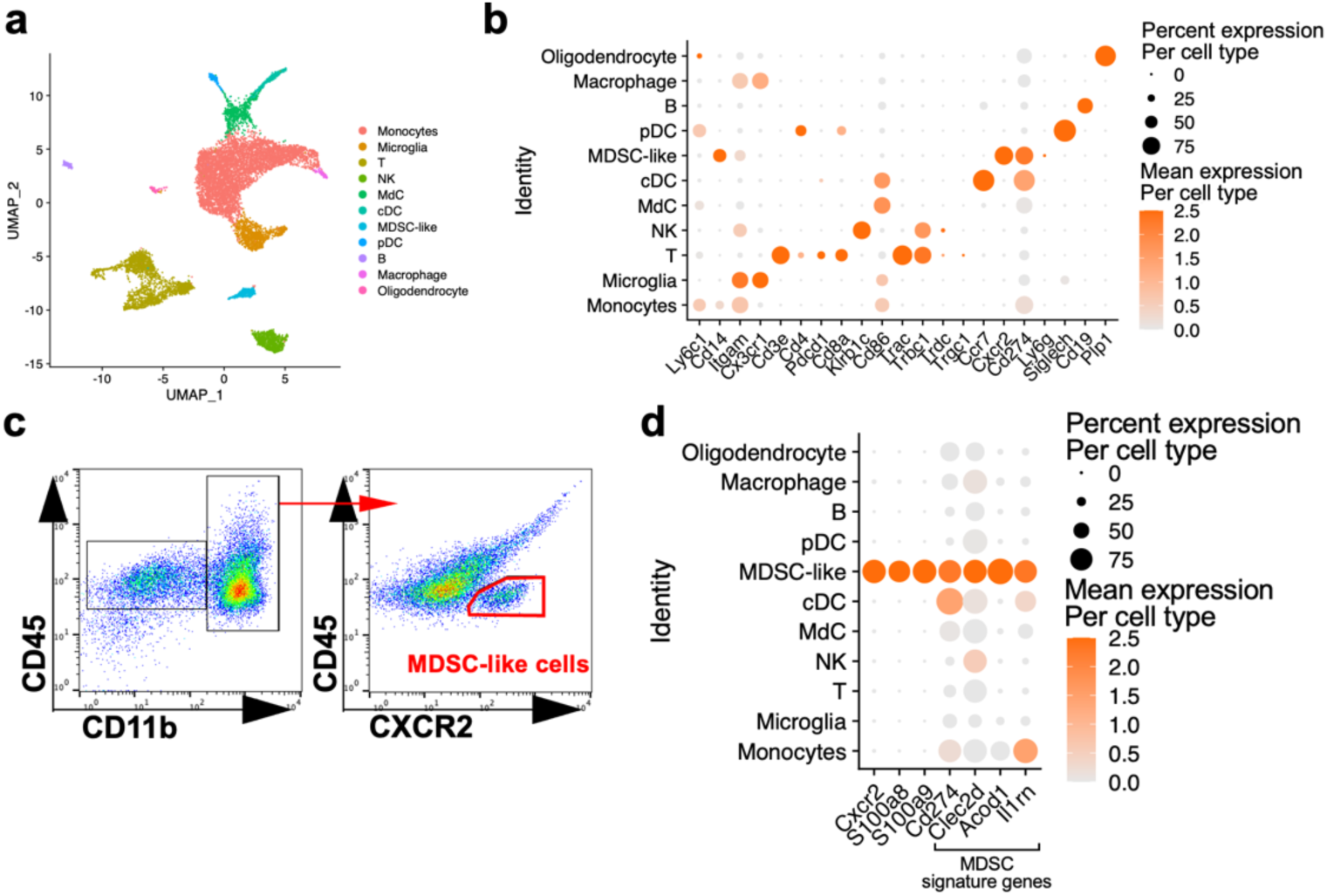
Identities of each immune cell in scRNA-seq, and the gating strategy used to identify CXCR2^+^ MDSC-like cells by flow cytometry, related to Fig. 2. (**a**) UMAP of CD45^+^ immune cell types in the OB at 6 dpi, clustered by scRNA-seq analysis. (**b**) Dot plot showing expression levels of cell type-specific marker genes within each immune cell cluster. (**c**) Representative flow cytometry plot showing the percentage of CXCR2^+^ MDSC-like cells among CD45^+^ CD11b^+^ myeloid cell population in the OB. (**d**) Dot plot showing expression of signature genes by MDSCs within each cell cluster.

**Supplementary Fig 3.**
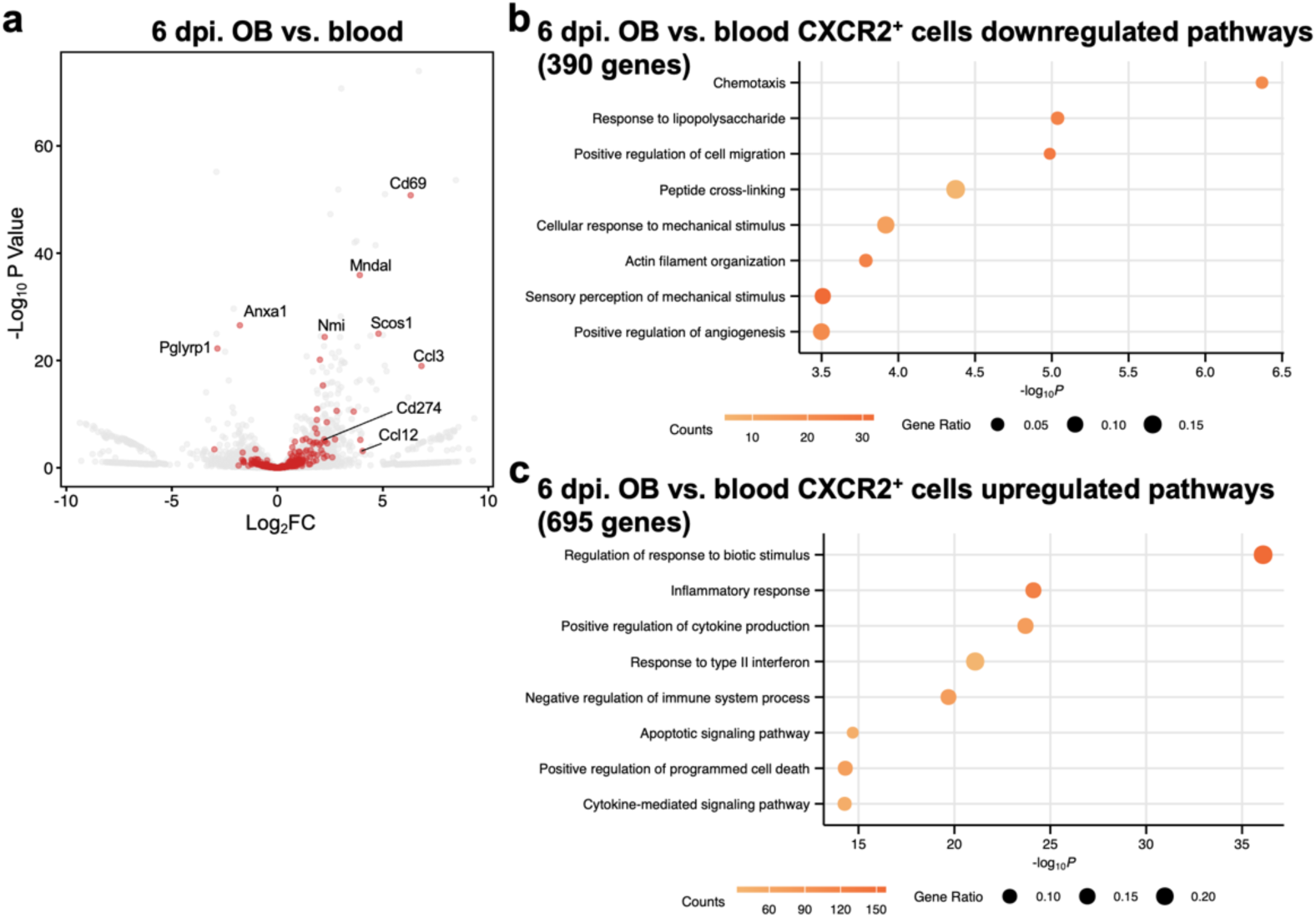
Genes differentially expressed by CXCR2^+^ cells in the OB and blood at 6 dpi, related to Fig. 4. (**a**) Volcano plot showing genes differentially expressed by CXCR2^+^ cells in the OB and blood (data from RNA-seq analysis). Gene ontology (GO) terms “negative regulation of immune system process” are highlighted. (**b,c**) The top downregulated and upregulated GO Biological Process terms for genes differentially expressed between CXCR2^+^ cells in the OB and blood (data from RNA-seq analysis). The number and ratio of each gene set is indicated.

**Supplementary Fig 4.**
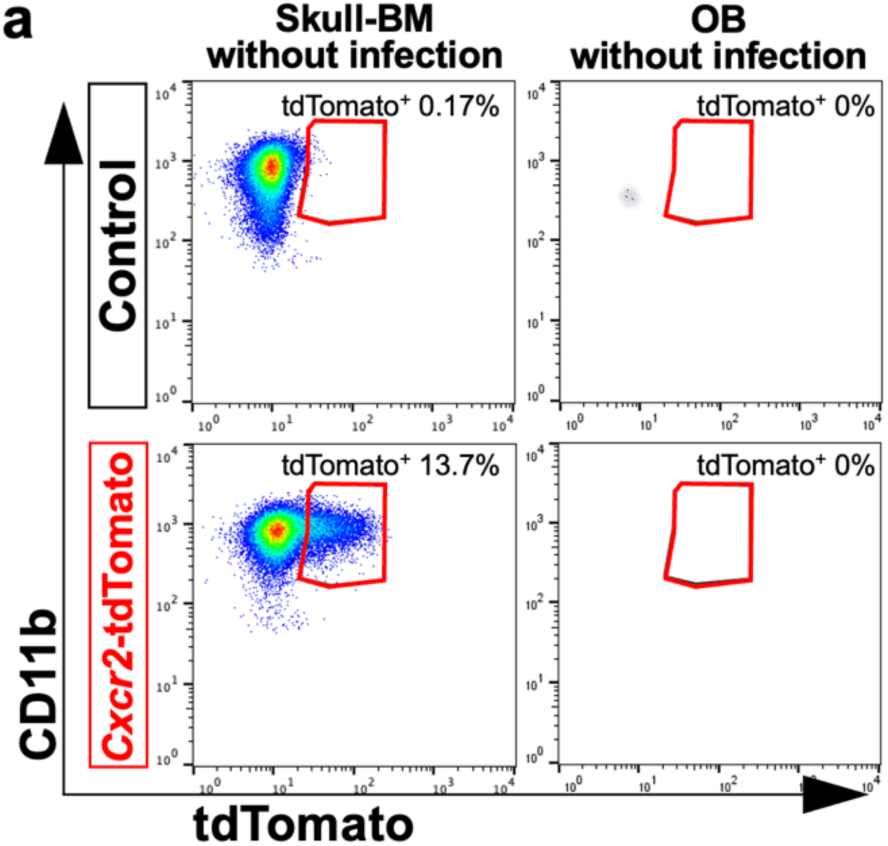
Percentage of tdTomato^+^ cells in the OB and skull-BM after tamoxifen (TAM) treatment in the absence of VSV infection, related to Fig. 4. (**a**) Representative flow cytometry plots showing the percentage of tdTomato^+^ cells among CD45^+^ CD11b^+^ CXCR2^+^ cells in the skull-BM and OB of TAM-treated *Cxcr2*-CreERT2 (control) mice (*n* = 1 mice) and TAM-treated *Cxcr2*-CreERT2; *Rosa*-tdTomato (*Cxcr2*-tdTomato) mice (*n* = 2 mice) in the absence of VSV infection.

**Supplementary Fig 5.**
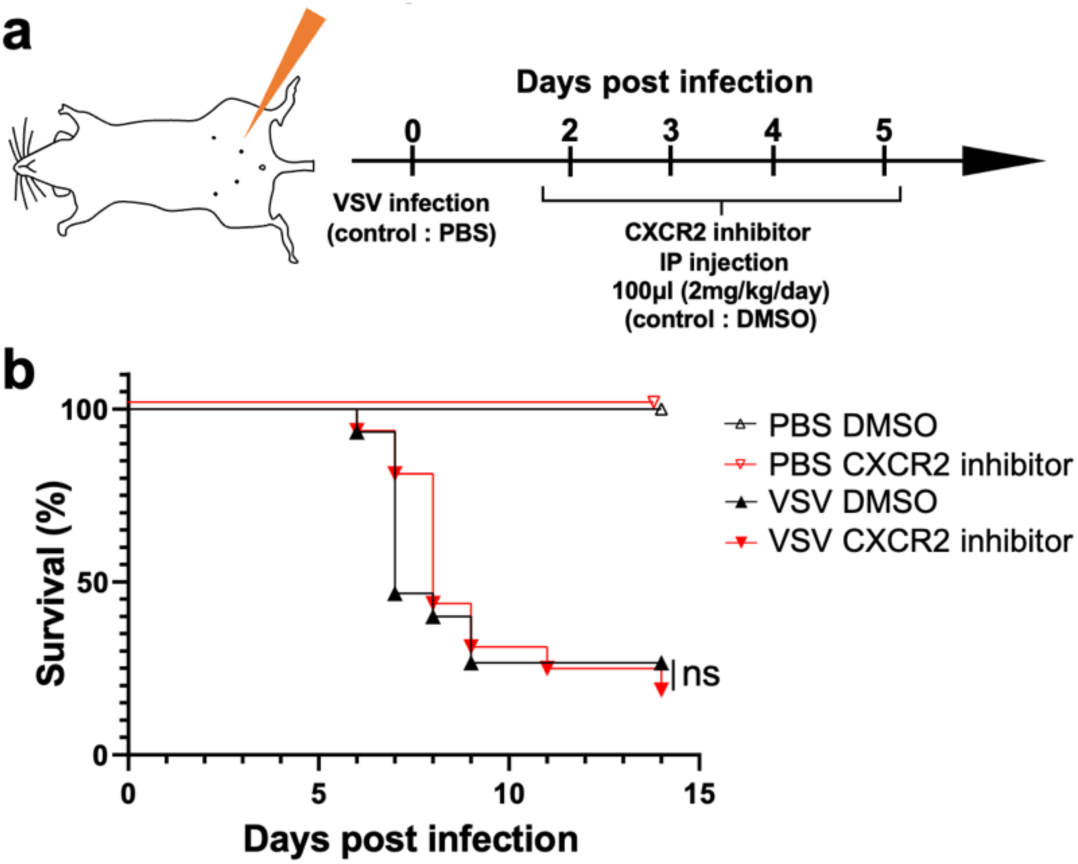
Intraperitoneal (IP) injection of a CXCR2 inhibitor does not alter survival rates after viral infection, related to Fig. 5. (**a**) General scheme of CXCR2 inhibitor injection. Briefly, a CXCR2 inhibitor in 100 μl DMSO was injected IP (2 mg/kg/day) during 2–5 dpi. (**b**) Survival curves for PBS control mice (*n* = 2 mice), PBS CXCR2 inhibitor-treated mice (*n* = 2 mice), VSV-infected control mice (*n* = 15 mice), and VSV-infected CXCR2 inhibitor-treated mice (*n* = 16 mice). ns, not significant. Statistical analyses were performed using log-rank test (b).

**Supplementary Fig 6.**
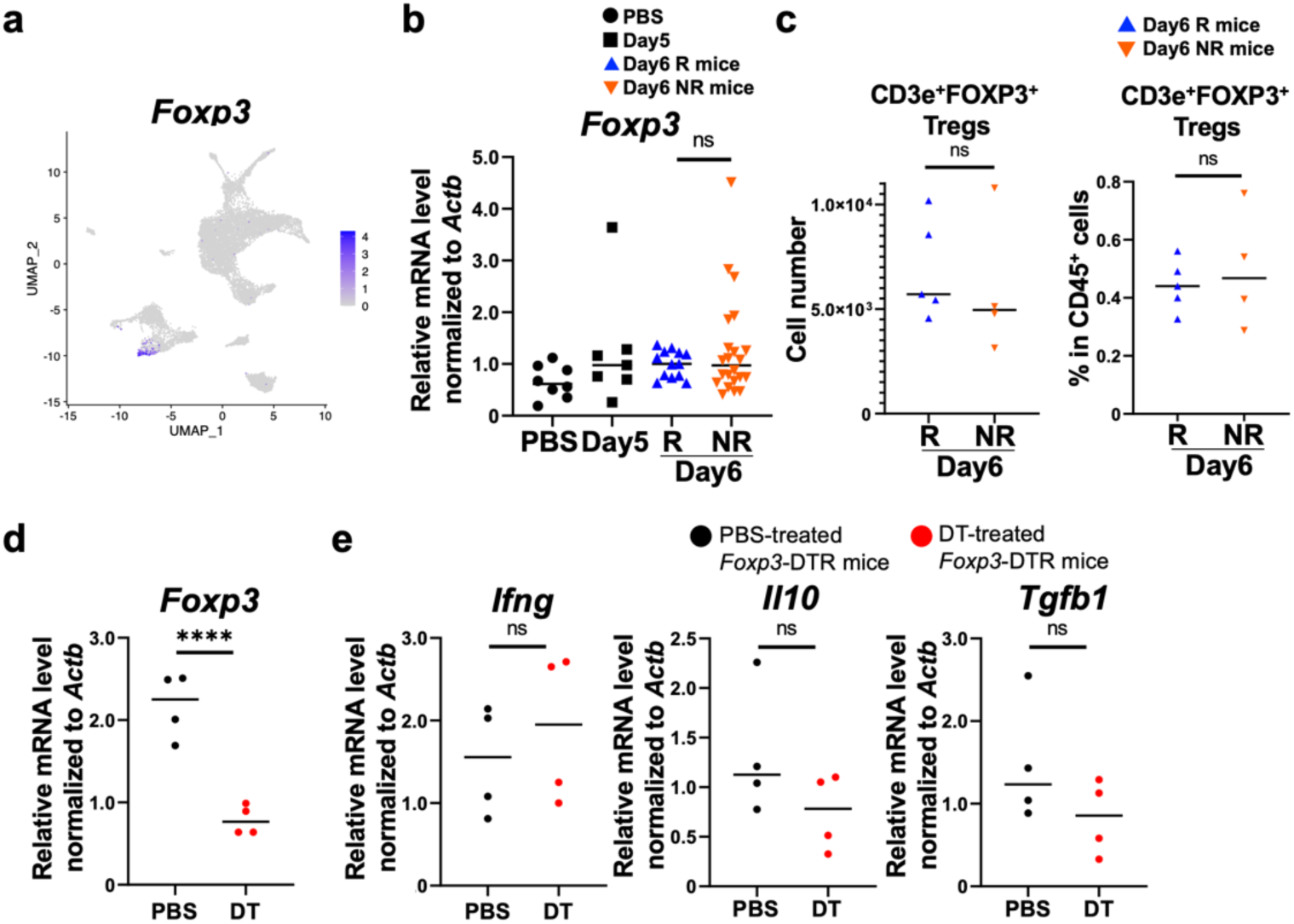
Regulatory T cells (Treg) do not expand following infection and have minimal impact on inflammation upon depletion. (**a**) Feature plot showing expression of *Foxp3* in the OB (from scRNA-seq analysis). (**b**) mRNA levels of *Foxp3* in the OB of uninfected (PBS) mice (*n* = 8 mice), VSV-infected (Day5) mice at 5 dpi (*n* = 7 mice), R mice (*n* = 13 mice), and NR mice (*n* = 22 mice). (**c**) Absolute number and percentage of FOXP3^+^ Tregs within the CD45^+^ immune cell population in the OB of R mice (*n* = 5 mice) and NR mice (*n* = 4 mice) at 6 dpi. (**d,e**) mRNA levels of *Foxp3*, *Ifng*, *Il10*, and *Tgfb1* in the OB of diphtheria toxin (DT)-treated *Foxp3*-diphtheria toxin receptor (DTR) mice (*n* = 4 mice) and PBS-treated *Foxp3*-DTR mice (*n* = 4 mice) at 6 dpi. DT (diluted in PBS, 25 ug/kg) was injected by IP injection every other day from one day before VSV infection. Data are presented as mean (b,c,d,e). ****p < 0.001, ns, not significant. Statistical analyses were performed using one-way ANOVA with Tukey’s post-tests (b) or unpaired two-sided Student’s t-tests (c,d,e).

**Supplementary Fig 7.**
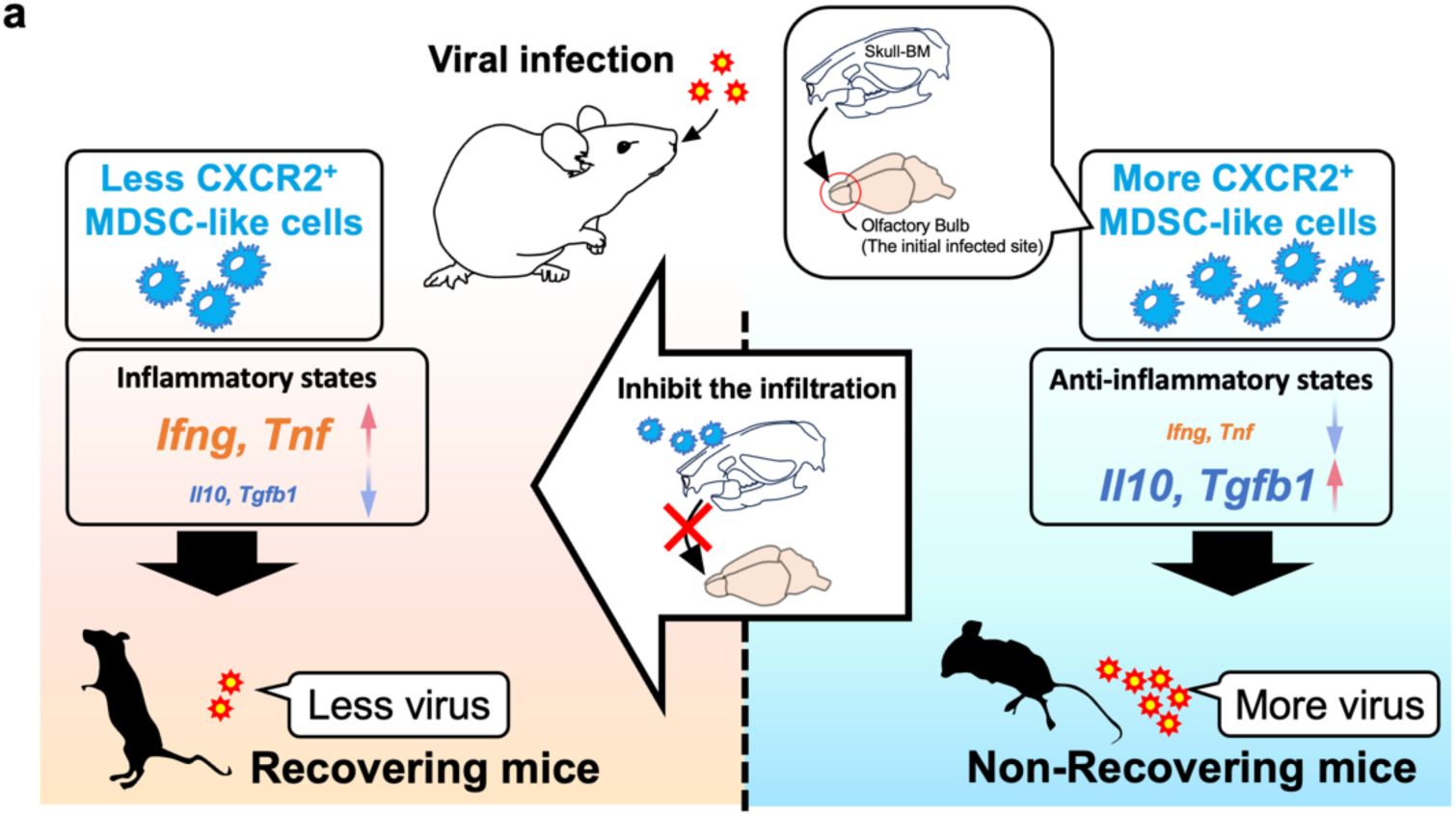
Local niche-derived immunosuppressive CXCR2^+^ cells impair antiviral immunity. (**a**) Schematic summary of the findings reported in this study.

**Supplementary Fig 8.**
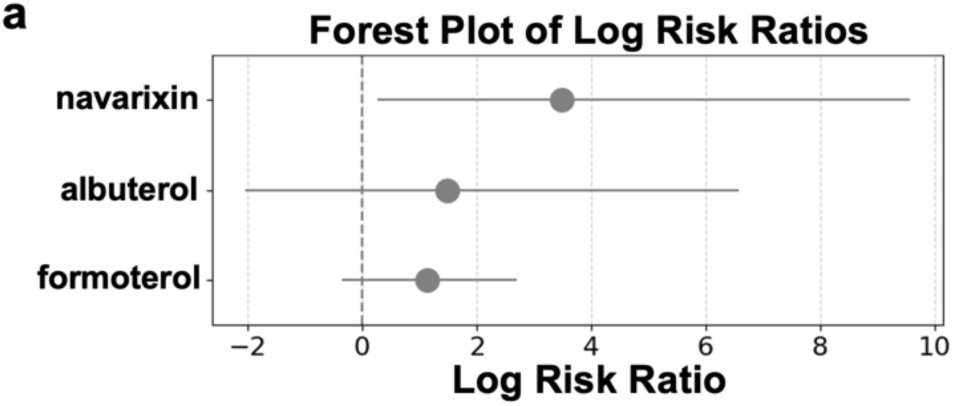
Inflammatory effects of navarixin in clinical trials. (**a**) Log risk ratio and 95% confidence interval (CI) values were calculated to investigate the inflammatory effects of navarixin. A difference was considered significant if the lower boundary of the 95% CI for the Log risk ratio was > 0. “Nasopharyngitis” was selected from the terms categorized under “Infections and infestations” in the organ_system category, as it was the only term commonly observed in clinical trials of the control drugs (albuterol and formoterol), which have similar clinical use similar to that of navarixin.

